# Social Isolation Causes Cortical and Trabecular Bone Loss in Adult Male, but not Female, C57BL/6J Mice

**DOI:** 10.1101/2023.01.27.525939

**Authors:** Rebecca V. Mountain, Audrie L. Langlais, Dorothy Hu, Roland Baron, Christine W. Lary, Katherine J. Motyl

## Abstract

Social isolation is a potent form of psychosocial stress and is a growing public health concern, particularly among older adults. Even prior to the onset of the COVID-19 pandemic, which has significantly increased the prevalence of isolation and loneliness, researchers have been concerned about a rising “epidemic” of loneliness. Isolation is associated with an increased risk for many physical and mental health disorders and increased overall mortality risk. In addition to social isolation, older adults are also at greater risk for osteoporosis and related fractures. While researchers have investigated the negative effects of other forms of psychosocial stress on bone, including depression and PTSD, the effects of social isolation on bone have not been thoroughly investigated. The aim of this study was to test the hypothesis that social isolation would lead to bone loss in male and female C57BL/6J mice. 16-week-old mice were randomized into social isolation (1 mouse/cage) or grouped housing (4 mice/cage) for four weeks (N=16/group). Social isolation significantly decreased trabecular (BV/TV, BMD, Tb. N., Tb. Th.) and cortical bone (Ct.Th., Ct.Ar., Ct.Ar./Tt.Ar., pMOI, Ct.Por.) parameters in male, but not female mice. Isolated male mice had signs of reduced bone remodeling represented by reduced osteoblast numbers, osteoblast-related gene expression and osteoclast-related gene expression. However, isolated females had increased bone resorption-related gene expression, without any change in bone mass. Overall, our data suggest that social isolation has negative effects on bone in males, but not females, although females showed suggestive effects on bone resorption. These results provide critical insight into the effects of isolation on bone and have key clinical implications as we grapple with the long-term health impacts of the rise in social isolation related to the COVID-19 pandemic.

## 1. Introduction

Social isolation and loneliness have gained recognition as major public health problems in light of the COVID-19 pandemic, particularly among older (60+) adults^1–3^. Social isolation refers to an objective lack of social interactions and connectedness, while loneliness is the subjective feeling of being isolated or alone^4^. Even prior to the pandemic, isolation and loneliness were described as a “behavioral epidemic”^5^, due to their widespread prevalence across the globe, affecting over one in four individuals over the age of 65^6^. Both social isolation and loneliness are associated with increased risk for numerous mental and physiological health conditions, including depression^7^, heart disease^8^, and dementia^9,10^. Additionally, social isolation has been identified to increase mortality risk by as much as 70%^11^, outweighing risk from high blood pressure, but similar to that of chronic smoking^12^.

While social isolation can affect individuals of any age, older adults are particularly vulnerable as they are more likely to live alone, have chronic health conditions that limit social interaction, or have suffered a loss of a partner or loved one^6^. Older adults are also the most at risk for osteoporosis, osteopenia, and related fractures. One in three women and one in five men over the age of 50 are expected to experience an osteoporosis-related fracture^13^. Despite this overlap in risk, there has been little research into the effects of social isolation on bone health. Previous studies have examined the effects of other forms of psychosocial stress on bone in rodents, including chronic subordinate colony housing^14,15^, early life stress^16^, and chronic mild stress^17,18^. These studies demonstrated psychosocial stress has negative effects on bone health. However, despite being a major public health concern, only five studies^19–23^ to our knowledge have directly,^19,21^ or indirectly,^20,22,23^ examined the effects of isolation on bone. Overall the limited research suggests that social isolation may negatively impact bone health, leading to decreased bone mineral density (BMD)^20,22,23^, although the findings are mixed^19,21^. None of these studies, however, have systematically investigated the effects of social isolation on bone metabolism or how it may differentially affect males and females.

The aim of this study was to test the effect of social isolation on bone metabolism in a rodent model. We further aimed to characterize how these effects may vary by sex and identify future mechanistic targets. We hypothesized that a four-week social isolation treatment would lead to reduced bone mineral density and trabecular microarchitecture in both males and females. This study is the first to characterize the effects of social isolation on bone metabolism in detail, and to investigate sex-specific effects of social isolation on bone. Our results will lead to future mechanistic and epidemiological studies examining the effects of social isolation on bone and may help identify individuals at higher risk for osteoporosis and related fractures.

## 2. Material and Methods

### 2.1 Mice

10-week-old female and male C57BL/6J mice were obtained from the Jackson Laboratory (Strain #000664, Bar Harbor, ME). Mice were acclimated in the barrier animal facility at MaineHealth Institute for Research (MHIR), an Association for Assessment and Accreditation of Laboratory Animal Care (AAALAC) accredited facility, for 6 weeks to ensure there were no residual effects of stress from transport. Mice were kept on a 14 hr light/10 hr dark cycle at room temperature (22°C) and given water and regular chow (Teklad global 18% protein diet, #2918, Envigo, Indianapolis, IN, USA) *ad libitum*. All procedures described in this study were approved by the MaineHealth Institute for Research Institutional Animal Care and Use Committee (IACUC).

### 2.2 Social Isolation

Two separate cohorts of 16-week old mice were included in this study. In the first cohort, mice were randomized into either socially isolated or control (grouped) housing by sex for four weeks (N=8/group). Isolated housing consisted of a single mouse per cage, with identical enrichment, in the form of a shepherd shack, to the control mice. Control mice were grouped housed, with four mice per cage, along with a shepherd shack. Isolated and grouped cages were interspersed within the same shelf to avoid any effects of differential shelf or row placement. This experimental procedure was repeated in a second cohort approximately two months later (N=8/group). Two female mice (one isolated, one grouped) in the second cohort were euthanized before the end of the treatment due to dermatitis and malocclusion.

### 2.3 Dual-energy x-ray absorptiometry (DXA)

Dual energy x-ray absorptiometry (DXA) was performed on both cohorts (N=16/group) using a Hologic Faxitron UltraFocus DXA system. The Faxitron was calibrated with bone and fat phantoms from the manufacturer before each scanning session. After weighing each mouse, total body (excluding the head) bone mineral density (aBMD), bone mineral content (aBMC), lean mass, and fat mass were measured, in addition to femoral and vertebral (L5) aBMD and aBMC. DXA was performed at baseline (16 weeks of age) to confirm there were no significant mean differences in bone or body parameters between isolated and grouped mice, and performed again at endpoint (20 weeks of age).

### 2.4 Micro-computed Tomography (μCT)

After sacrifice, femurs and L5 vertebrae were dissected and fixed in formalin (10% Neutral buffered) for 48 hours, and then transferred to 70% ethanol. X-ray micro computed tomography (μCT) was performed using a Scanco VivaCT-40 μCT system (Bassersdorf, Switzerland) with a resolution of 10.5 μm. The VivaCT-40 was used to assess cortical bone morphology of the femoral mid-diaphysis, and trabecular architecture of the distal femoral metaphysis in females from the first experimental cohort (N=8/group) and males from both cohorts (N=16/group). It was also used to assess trabecular architecture in the L5 vertebral body in the first cohort of both sexes (N=8/group). Bones were scanned in a 4-bone insert, all of which contained a sample during each scan. Scans were performed using an isotropic voxel size of 10.5 μm^3^, a 114 mA x-ray tube current with a 70 kVp peak x-ray tube intensity, and 250 ms integration time. Scans were then subjected to Gaussian filtration and segmentation and analyzed using Scanco analysis software.

### 2.5 Bone Turnover Markers

Blood was collected from all mice after decapitation and allowed to clot for at least 10 minutes. Samples were then centrifuged at 10,000 rpm for ten minutes at room temperature. Serum was then isolated, aliquoted, and stored at −80°C. Serum P1NP and CTX were measured in the first cohort (N = 4-7/group) using the Rat/Mouse P1NP enzyme and the RatLabs CTX-I immunoassays (EIA, Immunodiagnostic Systems, Boldon, UK). Assays were performed following the manufacturer’s instructions and read on a FlexStation 3 plate reader (Molecular Devices, Eugene, OR, USA). Results were calculated using a 4-parameter logistic curve.

### 2.6 Histomorphometry

Histomorphometric analyses were performed on the femur of male mice collected from the second cohort (N=7/group). Mice were injected with calcein (20 mg/kg, Sigma) 7 and 2 days before euthanasia for dynamic labeling measurements (N=4-6/group). Femurs were dissected and fixed in formalin (10%) for 48 hours, and transferred to 70% Dehydrant Alcohol. Femurs were dehydrated with graded acetone and embedded in methyl methacrylate. Longitudinal 4 μm sections were cut consecutively with a microtome and stained with either Von Kossa, 2% Toluidine Blue, fluorescent labeling, or TRAP to evaluate osteoid and osteoblasts, dynamic mineralized bone, or osteoclasts respectively. After collecting the initial first set of three slides, the femurs were sectioned again, skipping 120 microns, to collect an additional three slides, one for each of the fluorescent labeling, Toluidine Blue, and TRAP. The data from both sets (two slides/staining/sample, a total six slides) were then averaged together for the structure data (E.g., BV/TV, Tb.Th.). The cellular data (E.g., N. OB., N. OC.) and dynamic data (E.g., MAR, BFR) were averaged from two slides, either Toluidine Blue, fluorescent labeling, or TRAP stained.

Von Kossa slides were imaged with a Nikon E800 microscope and Olympus DP71 digital camera using Olympus CellSens software. Histomorphometric measurements were taken in a 1.3 mm x 0.9 mm region 225 μm away from the growth plate under 200-400X magnification and quantified using OsteoMeasure software (Osteometrics Inc., Decatur, GA, USA).

### 2.7 RNA Isolation and Real-time qPCR (RT-qPCR)

At the end of the four weeks of treatment, whole tibia were collected from the first cohort and flash frozen in liquid nitrogen before being stored at −80°C (N=8/group). RNA extraction was performed under liquid nitrogen conditions by crushing the bone and homogenizing in 1 mL of trizol. Samples were then incubated in 200 μL of chloroform for 10 minutes, and centrifuged. The aqueous layer was removed and 500 μL of isopropanol was added and frozen at −80°C overnight. Samples were centrifuged, the supernatant removed, and 1 mL of 75% ethanol added to wash RNA pellet twice. Supernatant was removed again and pellet allowed to air dry for 5 minutes before dissolving in nuclease-free water and freezing overnight at −80°C. RNA concentration was measured using NanoDrop 2000, and diluted appropriately (if >1000 ng/μL) with nuclease-free water based on concentration. RNA concentration was used to calculate appropriate cDNA mix using 10x RT buffer, 25x dNTP, 10x random primers, reverse transcriptase, and nuclease-free H2O. Samples were run on a PCR protocol of 10 minutes at 25°C, 120 minutes at 37°C, 5 minutes at 85°C, and cooled to 4°C. Samples were then diluted with 180 μL of nuclease-free water. 3 μL of cDNA were used for RT-qPCR analysis combined with nuclease-free water, SYBER green (BioRad, Hercules, CA), and appropriate forward and reverse primers. A BioRad® Laboratories CFX 384 real-time PCR system was used to analyze mRNA expression. Primers were purchased from Integrated DNA Technologies (IDT) (Coralvile, IA) or Qiagen (Germantown, MD), and primer sequences used in this study are listed in **Supplementary Table S1**. Hypoxanthine guanine phosphoribosyl transferase (*Hprt*) was used as the housekeeping gene for all analyses.

### 2.8 Statistical Analysis

All statistical analyses were performed using GraphPad Prism 9 XML Project® software. Data were analyzed using a Student’s t-test or two-way ANOVA, with α ≤ 0.05 considered statistically significant. Tukey’s *post hoc* test was performed if two-way ANOVA resulted in a significant interaction effect. Each cohort was initially examined separately, and data from both cohorts was combined for DXA and μCT of the male femurs only after no significant differences were found between the two cohorts.

## 3. Results

### 3.1 Social isolation caused bone loss in male, but not female, mice

Four weeks of social isolation decreased both trabecular and cortical bone parameters in the femur of male mice (**Figure 1**; **Tables 1 and 2**). Bone volume fraction (BV/TV) was reduced by 26% in isolated male mice compared to grouped mice, while bone mineral density (BMD) was reduced by 20%. Structural modeling index (SMI), trabecular number (Tb.N), thickness (Tb.Th), and spacing (Tb.Sp) were also significantly lower in the trabecular bone of isolated male mice relative to grouped mice. Isolated females had significantly lower Tb.Sp relative to grouped mice, but no other significant differences in trabecular bone.

**Figure 1.**
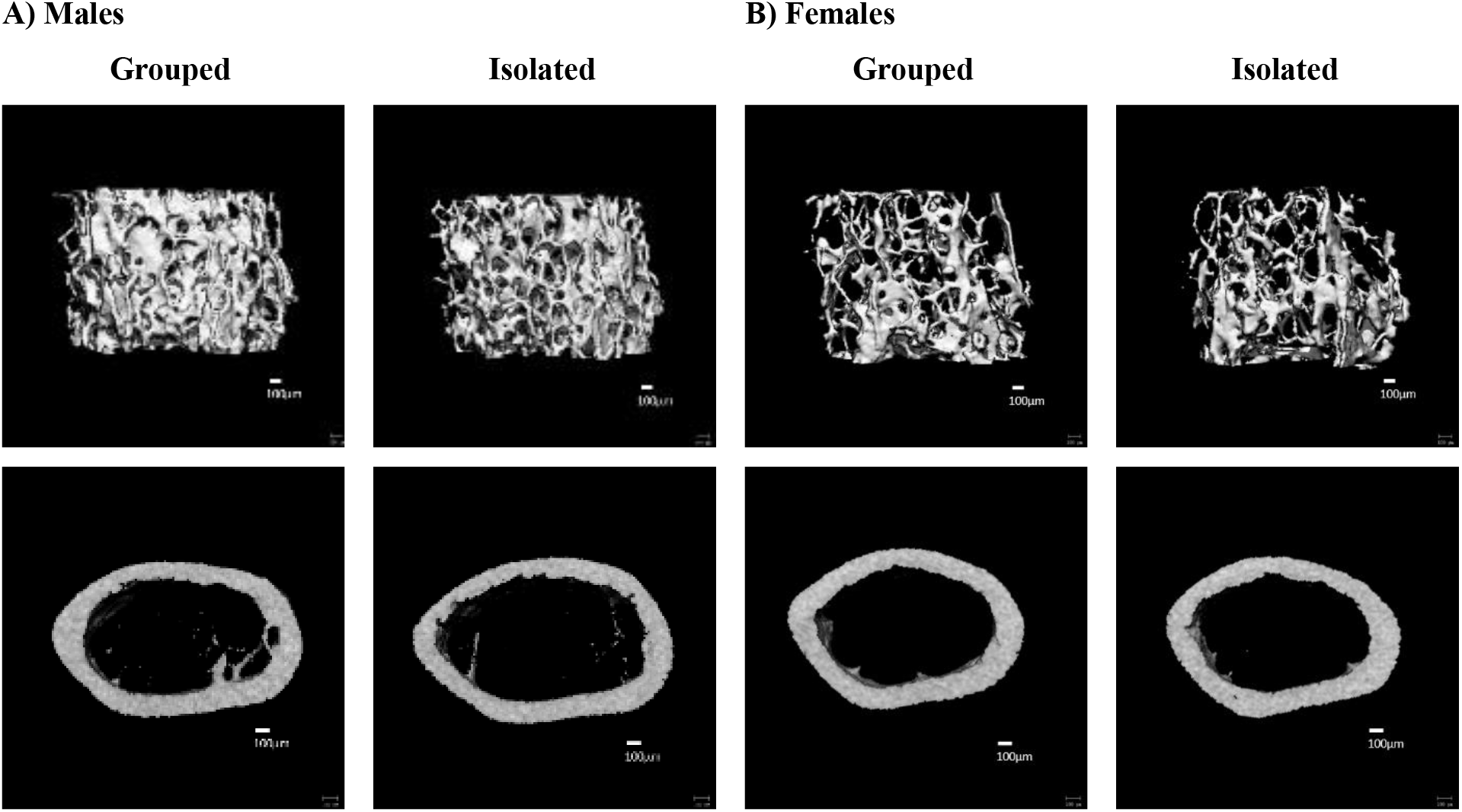
Social isolation decreased trabecular and cortical bone parameters in male mice, but did not affect bone parameters in females. 16-week old A) male and B) female mice were housed for four weeks in either social isolation (1 mouse/cage) or grouped housing (4 mice/cage). Changes in distal femur trabecular and cortical midshaft bone density and microarchitecture were measured using μCT. Scale bar = 100μm.

**Table 1.** Trabecular microarchitecture of distal femur of male and female mice housed in grouped or isolated housing for four weeks (Males: N=16/group Females: N=8/group).

|  | **Trabecular Bone** | **Grouped** | **Isolated** | **p-value** |
| --- | --- | --- | --- | --- |
| **Males** | BV/TV(%) | 21.9 ± 3.6 | 16.2 ± 2.1 | **<0.0001** |
|  | BMD (mg/cm<sup>3</sup>) | 222 ± 28 | 177 ± 16 | **<0.0001** |
|  | Conn.D (1/mm<sup>3</sup>) | 164 ± 21 | 154 ± 24 | 0.2221 |
|  | SMI | 1.51 ± 0.39 | 2.01 ± 0.24 | **0.0001** |
|  | Tb.N (1/mm) | 5.43 ± 0.30 | 5.13 ± 0.27 | **0.0065** |
|  | Tb.Th (mm) | 0.056 ± 0.004 | 0.048 ± 0.001 | **<0.0001** |
|  | Tb.Sp (mm) | 0.177 ± 0.011 | 0.187 ± 0.012 | **0.0113** |
| **Females** | BV/TV(%) | 8.2 ± 1.3 | 8.0 ± 1.6 | 0.8116 |
|  | BMD (mg/cm<sup>3</sup>) | 118 ± 9 | 118 ± 12 | 0.9693 |
|  | Conn.D (1/mm<sup>3</sup>) | 62 ± 13 | 52 ± 19 | 0.2124 |
|  | SMI | 2.77 ± 0.17 | 2.91 ± 0.32 | 0.2865 |
|  | Tb.N (1/mm) | 3.58 ± 0.21 | 3.52 ± 0.17 | 0.5693 |
|  | Tb.Th (mm) | 0.047 ± 0.002 | 0.050 ± 0.002 | **0.0245** |
|  | Tb.Sp (mm) | 0.279 ± 0.016 | 0.285 ± 0.016 | 0.4624 |

Data presented as mean ± SD.
BV/TV = bone volume/total volume; BMD = bone mineral density; Conn.D = connectivity density; SMI = structure model index; Tb.N = trabecular number; Tb.Th = trabecular thickness; Tb.Sp = trabecular separation.
Bold values indicate significance with $p < 0.05$ .

**Table 2.** Cortical microarchitecture of the femur midshaft of male and female mice housed in grouped or isolated housing for four weeks (Males: N=16/group; Females: N=8/group).

|  | **Cortical Bone** | **Grouped** | **Isolated** | **p-value** |
| --- | --- | --- | --- | --- |
| **Males** | Ct.Ar (mm<sup>2</sup>) | 0.926 ± 0.069 | 0.828 ± 0.058 | **0.0002** |
|  | Tt.Ar (mm<sup>2</sup>) | 2.09 ± 0.15 | 1.99 ± 0.17 | 0.0853 |
|  | Ma.Ar (mm<sup>2</sup>) | 1.17 ± 0.10 | 1.16 ± 0.13 | 0.9327 |
|  | Ct.Ar/Tt.Ar (%) | 44.3 ± 1.9 | 41.7 ± 2.2 | **0.0011** |
|  | Ct. TMD (mg/cm<sup>3</sup>) | 1260 ± 16 | 1253 ± 18 | 0.2088 |
|  | Ct.Por (%) | 0.416 ± 0.053 | 0.474 ± 0.096 | **0.0422** |
|  | Ct.Th (mm) | 0.187 ± 0.011 | 0.169 ± 0.009 | **<0.0001** |
|  | pMOI (mm<sup>4</sup>) | 0.515 ± 0.066 | 0.452 ± 0.067 | **0.0116** |
| **Females** | Ct.Ar (mm<sup>2</sup>) | 0.826 ± 0.027 | 0.828 ± 0.037 | 0.9304 |
|  | Tt.Ar (mm<sup>2</sup>) | 1.73 ± 0.06 | 1.78 ± 0.12 | 0.3163 |
|  | Ma.Ar (mm<sup>2</sup>) | 0.91 ± 0.04 | 0.95 ± 0.10 | 0.2479 |
|  | Ct.Ar/Tt.Ar (%) | 47.7 ± 0.8 | 46.6 ± 2.7 | 0.2916 |
|  | Ct. TMD (mg/cm<sup>3</sup>) | 1295 ± 15 | 1291 ± 13 | 0.5689 |
|  | Ct.Por (%) | 0.415 ± 0.030 | 0.442 ± 0.028 | 0.0857 |
|  | Ct.Th (mm) | 0.194 ± 0.006 | 0.192 ± 0.012 | 0.6425 |
|  | pMOI (mm<sup>4</sup>) | 0.369 ± 0.026 | 0.389 ± 0.046 | 0.3068 |

Data presented as mean ± SD.
Ct. = cortical; Ar. = area; Tt. = total; Ma. = medullary; Ct.Ar/Tt.Ar= cortical area/total area; Ct.TMD = cortical tissue mineral density; Ct.Por. = cortical porosity; Ct.Th. = cortical thickness; pMOI = polar moment of inertia.
Bold values indicate significance with $p < 0.05$ .

Male isolated mice also had significantly lower femoral cortical parameters. Cortical thickness (Ct.Th) was 9% lower in isolated male mice relative to grouped mice. Cortical area (Ct.Ar), cortical area fraction (Ct.Ar/Tt.Ar), and polar moment of inertia (pMOI) were also significantly lower than grouped-housed mice, while cortical porosity (Ct. Por.) was significantly increased. Female mice did not have any significant differences in cortical bone between groups.

A similar pattern was seen in the L5 vertebrae in males, where social isolation decreased vertebral BV/TV by 19%, and BMD by 16%. SMI and Tb.Th. were also significantly reduced in the L5 in males (**Table 3**). Socially isolated females did not have any significant differences in bone parameters in the L5.

**Table 3.** Trabecular microarchitecture of L5 vertebrae of male and female mice housed in grouped or isolated housing for four weeks (Males and females: N=8/group).

|  | **Trabecular Bone** | **Grouped** | **Isolated** | **p-value** |
| --- | --- | --- | --- | --- |
| **Males** | BV/TV(%) | 29.5 ± 2.5 | 23.8 ± 2.0 | **0.0002** |
|  | BMD (mg/cm<sup>3</sup>) | 299 ± 24 | 250 ± 15 | **0.0002** |
|  | Conn.D (1/mm<sup>3</sup>) | 249 ± 22 | 260 ± 27 | 0.3890 |
|  | SMI | 0.35 ± 0.20 | 0.74 ± 0.15 | **0.0006** |
|  | Tb.N (1/mm) | 5.64 ± 0.33 | 5.36 ± 0.34 | 0.1183 |
|  | Tb.Th (mm) | 0.054 ± 0.003 | 0.046 ± 0.001 | **<0.0001** |
|  | Tb.Sp (mm) | 0.170 ± 0.011 | 0.181 ± 0.012 | 0.1076 |
| **Females** | BV/TV(%) | 24.0 ± 2.8 | 22.4 ± 1.4 | 0.1730 |
|  | BMD (mg/cm<sup>3</sup>) | 257 ± 24 | 244 ± 11 | 0.1893 |
|  | Conn.D (1/mm<sup>3</sup>) | 140 ± 27 | 131 ± 20 | 0.4746 |
|  | SMI | 0.58 ± 0.22 | 0.67 ± 0.16 | 0.3598 |
|  | Tb.N (1/mm) | 4.10 ± 0.35 | 3.91 ± 0.23 | 0.2207 |
|  | Tb.Th (mm) | 0.057 ± 0.002 | 0.055 ± 0.001 | 0.1064 |
|  | Tb.Sp (mm) | 0.247 ± 0.022 | 0.261 ± 0.016 | 0.1813 |

Data presented as mean ± SD.
BV/TV = bone volume/total volume; BMD = bone mineral density; Conn.D = connectivity density; SMI = structure model index; Tb.N = trabecular number; Tb.Th = trabecular thickness; Tb.Sp = trabecular separation.
Bold values indicate significance with $p < 0.05$ .

There were no significant differences in weight or other tissue parameters in male mice observed via DXA. Isolated female mice, on the other hand, had decreased fat mass relative to grouped mice (**Supplementary Figure S1**).

### 3.2 Social isolation increased serum bone formation markers in females, but not males

In order to determine if the differences between groups seen in μCT were a result of changes in bone formation, resorption, or both, we first examined bone turnover markers in serum (**Figure 2**). There were no significant differences between isolated and grouped males in either P1NP, indicative of bone formation, or CTX, indicative of bone resorption. Isolated females, however, had significantly higher levels of P1NP, but no differences in CTX.

**Figure 2.**
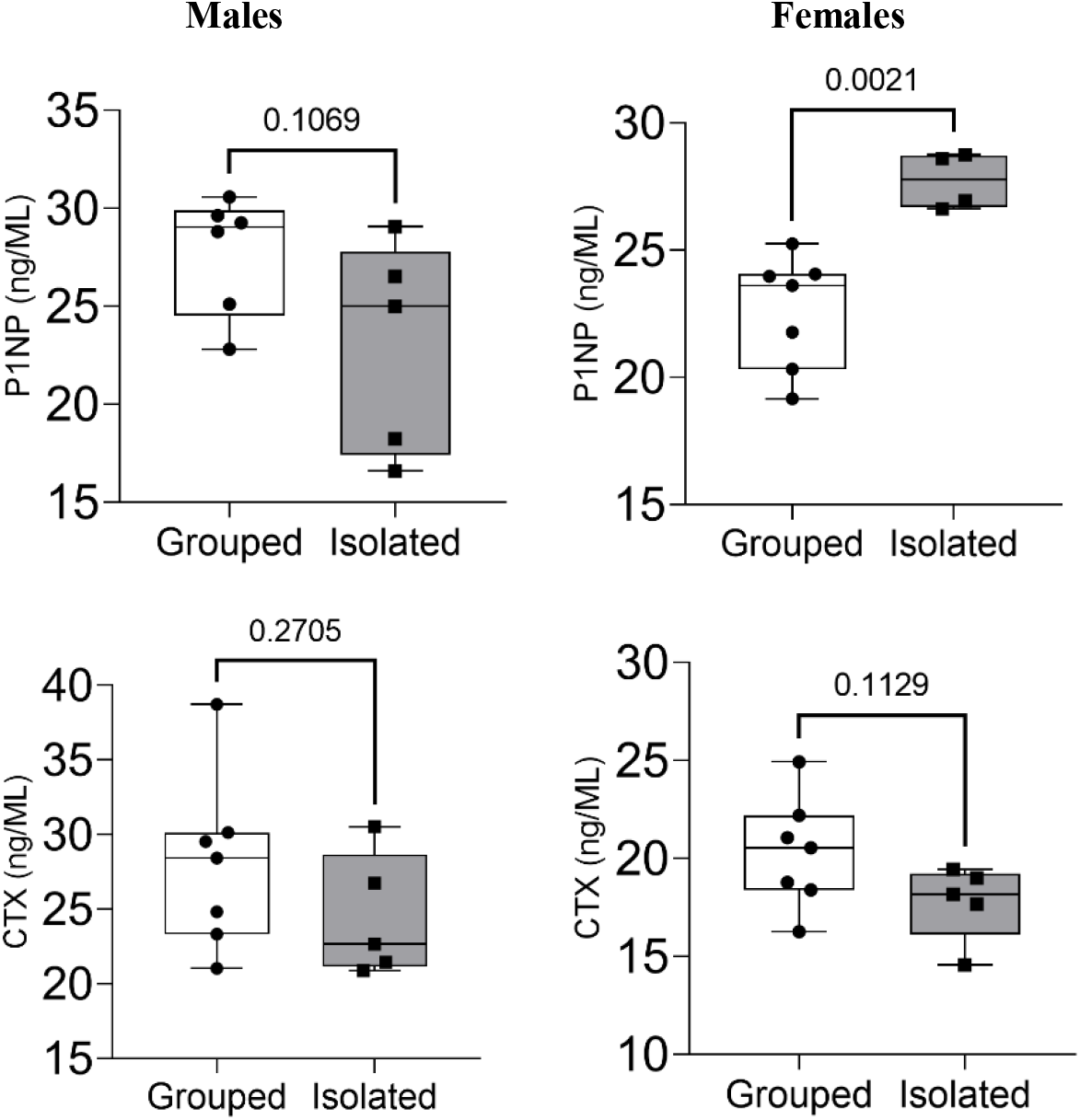
Social isolation significantly increased bone formation markers in females, but had no significant effect in males. Bone formation (P1NP) and resorption (CTX) markers were measured with serum EIA assays. N = 4-7/group.

### 3.2 Socially isolated male mice may have reduced osteoblasts, but not osteoclasts in the femur, relative to grouped mice

To further examine differences in bone turnover, we conducted histomorphometric analyses on femurs isolated from male mice (**Figure 3**). Consistent with μCT, social isolation reduced bone volume fraction, trabecular thickness, number, and spacing (**Table 4**). Additionally, the percent of trabecular bone surface covered by osteoblasts tended to be lower in the isolated mice (*p* = 0.07), potentially indicating a reduction in bone formation. There were no significant differences in osteoclasts, or in any of the dynamic labeling measurements.

**Figure 3.**
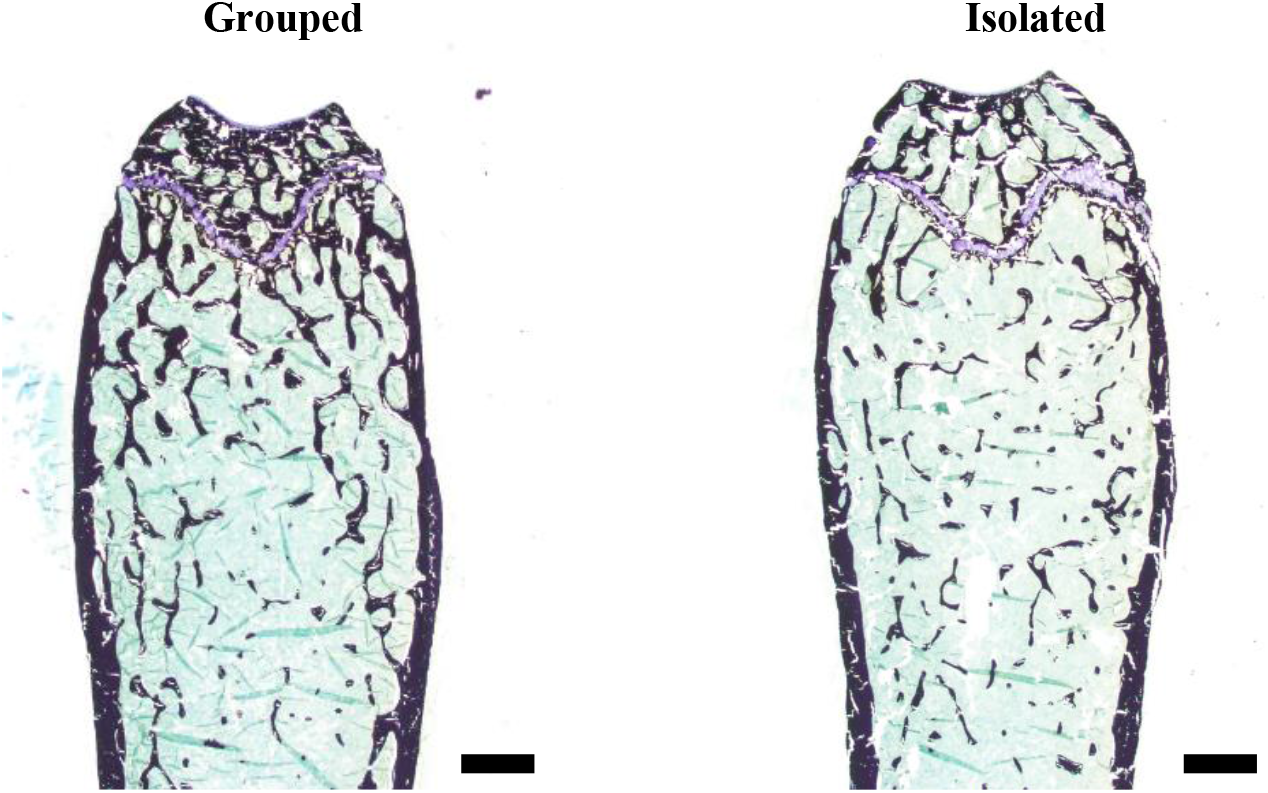
Social isolation decreased trabecular bone in male mice. Representative histomorphometry images of distal femur in male mice. Measures were taken in a 1.3 mm x 0.9 mm region 225 μm away from the growth plate. Scale bar = 500μm

**Table 4.** Trabecular histomorphometry of distal femur of male mice housed in grouped or isolated housing for four weeks. N=7/group. *Dynamic labeling measurements N=4-6/group.

|  | **Grouped** | **Isolated** | **p-value** |
| --- | --- | --- | --- |
| BV/TV (%) | 18.6 ± 4.2 | 13.6 ± 3.5 | **0.03** |
| Tb.Th (μm) | 48.3 ± 6.5 | 40.1 ± 6.2 | **0.03** |
| Tb.N (/mm) | 3.82 ± 0.42 | 3.32 ± 0.35 | **0.03** |
| Tb.Sp (μm) | 218 ± 33 | 267 ± 35 | **0.02** |
| MAR (μm/day)* | 1.15 ± 0.12 | 1.05 ± 0.13 | 0.24 |
| MS/BS (%)* | 49.38 ± 5.30 | 51.16 ± 7.04 | 0.66 |
| BFR/BV (%/day)* | 2.61 ± 0.41 | 2.68 ± 0.31 | 0.78 |
| BFR/BS (μm<sup>3</sup>/μm<sup>2</sup>/day)* | 0.57 ± 0.08 | 0.54 ± 0.12 | 0.65 |
| Ob.S/B.Pm (%) | 17.3 ± 3.8 | 12.7 ± 4.9 | 0.07 |
| N.Ob/B.Pm (#/mm) | 12.1 ± 2.2 | 9.2 ± 3.7 | 0.10 |
| OS/BS (%) | 10.0 ± 5.3 | 8.4 ± 4.6 | 0.57 |
| O.Th (μm) | 2.68 ± 0.17 | 2.52 ± 0.27 | 0.22 |
| Oc.S/BPm (%) | 9.80 ± 1.91 | 9.42 ± 1.40 | 0.68 |
| N.Oc/B.Pm (#/mm) | 4.75 ± 0.78 | 4.56 ± 0.63 | 0.62 |

Data presented as mean ± SD.
BV/TV = bone volume fraction; Tb.Th = trabecular thickness; Tb.N = trabecular number; Tb.Sp = trabecular separation; MAR = mineral apposition rate; MS/BS = mineralizing surface per unit bone surface; BFR/BV = bone formation rate per unit bone volume; BFR/BS = bone formation rate per unit bone surface; Ob.S/B.Pm = trabecular bone surface covered with osteoblasts; N.Ob/B.Pm = number of osteoblasts on bone surface; OS/BS = osteoid surface; O.Th = osteoid thickness; Oc.S/BPm = trabecular bone surface covered with osteoclasts; N.Oc/B.Pm = number of osteoclasts on bone surface.
Bold values indicate significance with $p < 0.05$ .

### 3.3 Social isolation had sex-specific effects on formation and resorption-related gene expression

Consistent with the trend toward reduced osteoblast numbers, isolated male mice had significantly decreased expression of *Runx2* and *Dmp1* in bone, indicating a decrease in osteoblast differentiation and bone mineralization (**Figure 4**). Social isolation, however, also caused significant decreases in gene expression related to bone resorption in males, specifically *Acp5* and *Ctsk*, relative to grouped housing. There were no differences in osteocalcin (*Bglap*), RANKL (*Tnfs11*), or osteoprotegerin (*Tnfsf11b*) between housing conditions in male mice. The ratio of RANKL to OPG (*Tnfs11/Tnfs11b*), however, tended to be lower in isolated males relative to grouped mice (*p* = 0.0548).

**Figure 4.**
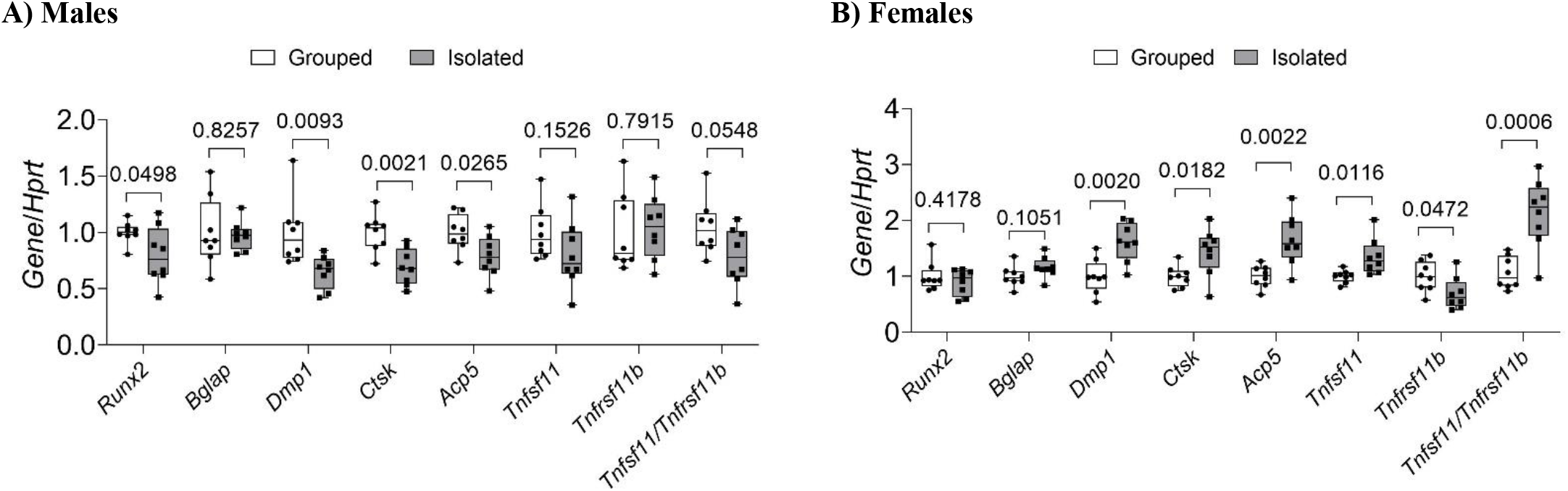
Social isolation decreased expression of bone formation and resorption marker genes in males and increased bone resorption marker gene expression in females in whole bone. Gene expression, normalized to *Hprt*, from whole tibia was measured in the first cohort of A) male and B) female 20 week-old mice isolated or grouped for four weeks using RT-qPCR. N=8/group.

Conversely, female isolated mice had a significant increase in expression of *Dmp1*. Isolated females also had significantly increased expression of genes, *Acp5, Ctsk*, and *Tnfs11*, related to bone resorption despite a lack of a structural bone phenotype. The ratio of *Tnfs11/Tnfs11b* was also significantly higher in isolated females relative to grouped females. There were no differences in osteocalcin (*Bglap*) or *Runx2* in females.

## 4. Discussion

The main finding of this study is that four weeks of social isolation led to significant reductions in both trabecular and cortical bone in male mice. Isolated male mice had a 26% and 19% reduction in bone volume fraction in the femur and L5 respectively, and a 9% reduction in femoral cortical thickness. These effect sizes are comparable to those seen four weeks after loss of sex hormones (OVX^24,25^ and ORX^26,27^), and half as large as those seen in hind limb unloading studies^28,29^. Social isolation, however, did not lead to bone loss in female mice. This is the first study, to our knowledge, to specifically test the effects of social isolation-induced bone loss through a range of imaging, genetic, and histologic approaches.

Our findings are consistent with Nagy et al.^22^, who found that 9 weeks of single housing reduced BMD in 3-week-old male C57BL/6J mice. Morey-Holton et al.^23^ also found that social isolation reduced bone formation in juvenile male rats, specifically during spaceflight. In contrast to our results, Tahimic et al.^20^ found that one month of social isolation reduced BV/TV and trabecular number, and increased trabecular spacing in adult female mice. Tahimic et al., however, used C57BL6/NJ mice in their experiments, and their results may indicate that sex differences in isolation-induced bone loss may be strain-specific. Additional experiments investigating inter-strain differences are needed to test this.

In our study, in addition to the overall trabecular and cortical bone loss, social isolation led to a reduction in both formation- and resorption-related gene expression in males. The quantity of bone surface covered in osteoblasts also trended lower in isolated males, but there were no differences in osteoclast numbers or in serum CTX. This indicates a low turnover phenotype in males, and also suggests the bone loss may be primarily driven by reduced bone formation, but this will need to be investigated further in our future studies. There were also no significant changes in weight or other body parameters in males, which suggests the differences between groups are unlikely due to differences in physical activity, fighting between grouped mice, diet, or other external factors.

Females, in contrast, had higher bone formation and resorption markers after exposure to social isolation, but the balance between the two was maintained since these changes did not have any significant effects on the bone phenotype aside from a decrease in trabecular spacing. These sex differences may be a result of the general protective effects of estrogen on bone^30^, or may suggest different mechanisms are at work within each sex. Previous studies investigating the effects of social isolation on other tissues, such as the brain, have also identified major sex differences^31^ which may operate through sex-specific mechanisms.

One mechanism by which social isolation stress may cause changes in bone in both sexes is through altered glucocorticoid signaling. Social isolation frequently increases circulating glucocorticoids, specifically cortisol in humans, and corticosterone in mice^32^. Glucocorticoid-induced osteoporosis, typically resulting from chronic use of pharmaceutical glucocorticoids, is the most common form of secondary osteoporosis^33^, and is characterized by a biphasic pattern of increased bone resorption and decreased formation followed by a prolonged period of decreased resorption and formation^34^. In this study, males showed a reduction in osteoblasts via histomorphometry, as well as osteoblast and osteoclast gene expression, in conjunction with bone loss. Females, however, showed an increase in both markers of bone formation and resorption without a change in bone phenotype. It is possible that these sex differences are due to a combination of the specific effects of estrogen on glucocorticoid signaling^35^ and the biphasic nature of glucocorticoid induced bone loss, leading to a delay in bone loss in females relative to males.

Another possible mechanism by which social isolation may be operating in bone is via elevated sympathetic nervous system (SNS) activity. Chronic stress activates the SNS, potentially increasing catecholamines, specifically norepinephrine and epinephrine^17,36^, although these findings have been less consistent among social isolation models^37,38^. If catecholamines are increased in response to social isolation, this could reduce bone mass via beta-2 adrenergic (β2AR) signaling^39,40^. Another possibility is that social isolation at room temperature (versus thermoneutral) leads to thermal stress in mice and consequently heightened SNS activity^41,42^ to stimulate brown adipose thermogenesis. This could then lead to bone loss due to cold stress^43–46^. To our knowledge, the effect of social isolation on cold stress and bone loss has not been explored and should be investigated in future studies.

This study had several major strengths. Our model of social isolation was designed to be clinically relevant to human social isolation, and therefore included enrichment in the form of shepherd shacks and did not eliminate other sensory input. Still, we saw dramatic bone loss in our male mice, suggesting even mild isolation has an effect on bone. By repeating this experiment in a second cohort we were also able to replicate our initial results in which social isolation led to bone loss in male mice, but not females. Additionally, we examined the effects of social isolation on bone in both sexes, which to our knowledge, had not been previously examined within the same study. Our study also examined the effects of social isolation on bone in both the axial and appendicular skeleton and at multiple levels, from μCT imaging and histomorphometry, to serum level turnover markers and mRNA expression. This thorough examination allowed us to identify changes in bone turnover between groups and provided insight into possible mediating cell types.

There were also several limitations to this study. First, our bone turnover assays, P1NP and CTX, included ≤5 mice per group, and were likely underpowered. The dynamic labeling was also not successful in all mice injected with calcein, and therefore only 4-6 mice/group were able to be analyzed for dynamic histomorphometry. Second, we were not able to evaluate the effects of social isolation on stress-related depressive or anxious behavior in the mice, which could provide better insight into the effects of social isolation stress in each sex and the resulting effects on bone. We also were unable to perform metabolic cage studies on grouped mice, and thereby identifying how housing conditions affected activity levels. Lastly, while the results of this study suggest the bone loss seen in males may be primarily related to decreased bone formation, specific mediating mechanisms were not explored and will be the subject of future analyses.

Future studies will focus on further investigating the sex differences in social isolation-induced bone loss and mediating pathways including glucocorticoid and adrenergic signaling. It is possible that females would experience bone loss with prolonged exposure to social isolation. A time-course experiment would help elucidate the sex-specific effects, as well as the effects of social isolation on different stages of bone formation and resorption in both sexes.

Overall, the findings of this study demonstrate that four weeks of social isolation reduces cortical and trabecular bone in male adult mice, but not females. Future studies should focus on potential mediating mechanisms and determine if the sex differences observed in this study are a result of sex-specific mechanisms. Collectively, the results of this project shed critical new light on the negative effects of social isolation on skeletal health, and have key clinical implications as we grapple with the long-term health impacts of the rise in social isolation related to the COVID-19 pandemic. These results will also lead to future mechanistic and epidemiological studies, which will improve our ability to identify at-risk individuals and possible treatments in the future.

## Acknowledgments

The authors thank Patrizia Roy for technical assistance. Funding: This work was supported by the NIH National Institute of Arthritis and Musculoskeletal and Skin Diseases (R01AR076349 to KJM). This work was also supported by the National Institute of General Medical Sciences through the Northern New England Clinical and Translational Research (NNE-CTR) Network (U54GM115516), the MaineHealth COBRE in Mesenchymal and Neural Regulation of Metabolic Networks (P20GM121301).

## Supplementary Materials

**Supplementary Table S1.**
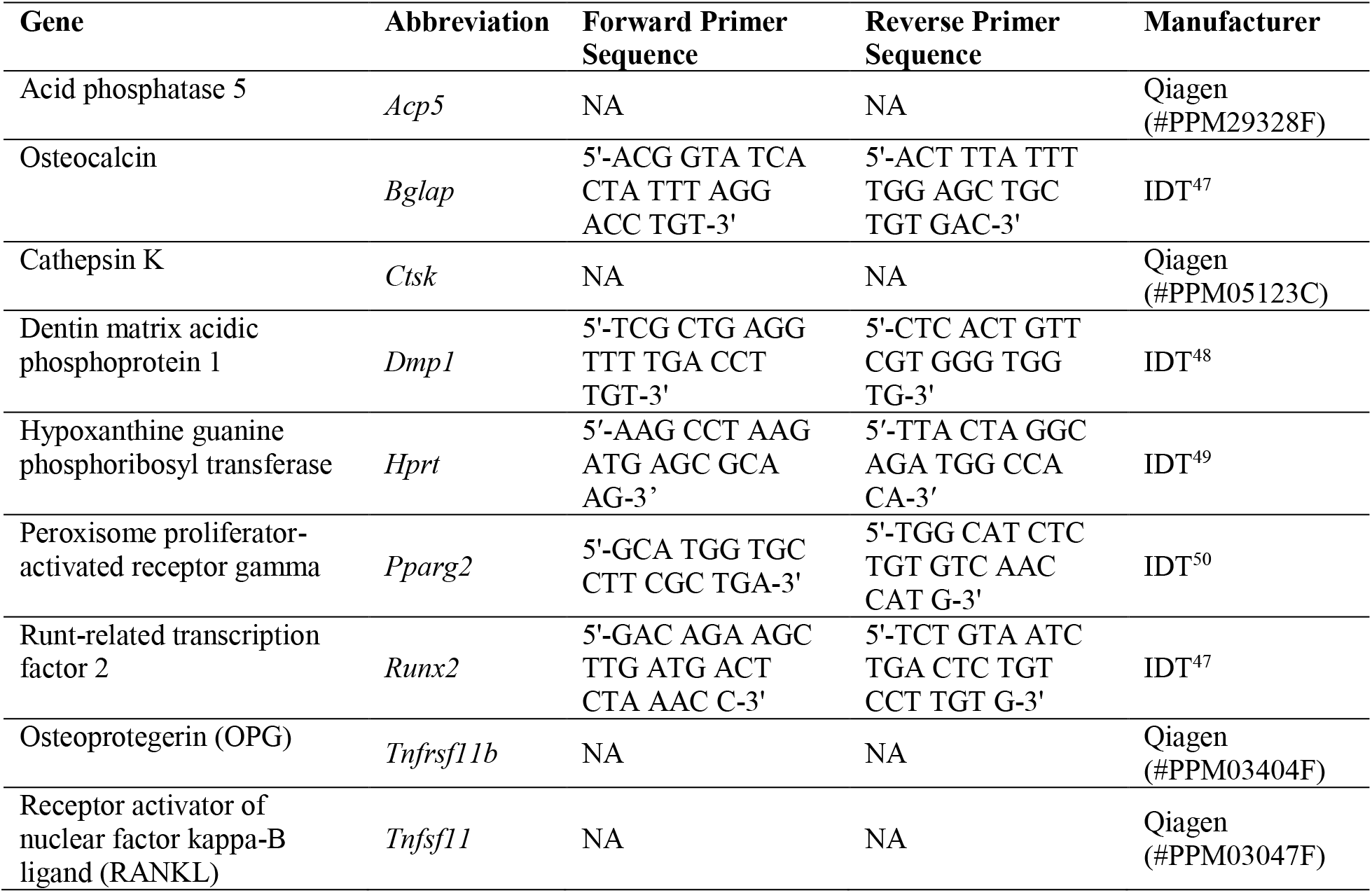
Primer sequences used for qPCR.

**Supplemental Figure S1.**
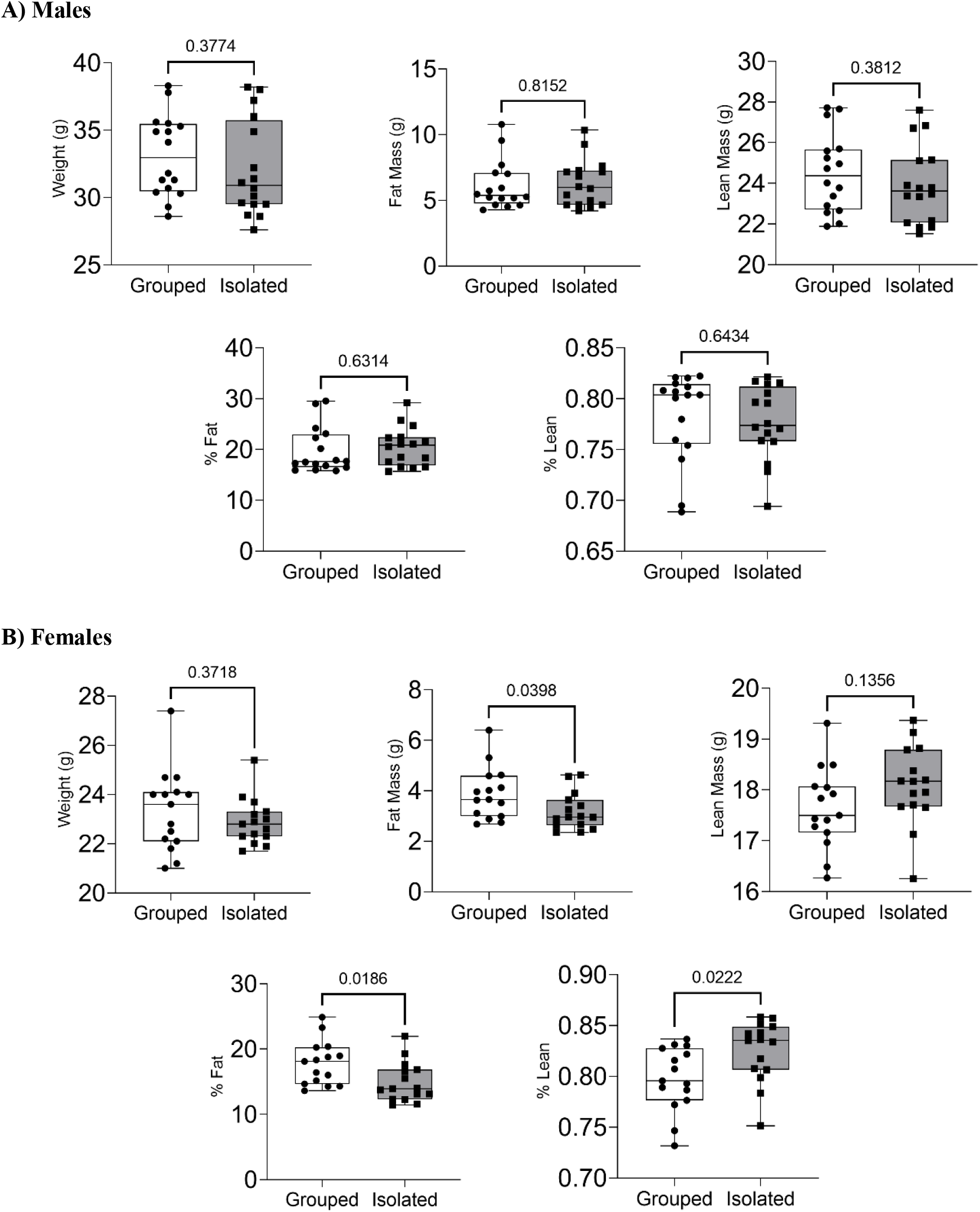
Social isolation decreased fat mass in female mice, but did not significantly affect body parameters in male mice. At the end of 4 weeks of treatment, male and female mice body parameters were measured using dual-energy x-ray absorptiometry (DXA). N=16/group.

## Notes

**Declarations of Interest:** None

### Competing Interest Statement

The authors have declared no competing interest.

